# Bluetongue virus serotype 12 in the Netherlands in 2024 - A BTV serotype reported in Europe for the first time

**DOI:** 10.1101/2024.12.03.626362

**Authors:** René van den Brom, Inge Santman-Berends, Mark G. van der Heijden, Frank Harders, Marc Engelsma, Rene G.P. van Gennip, Mieke A. Maris-Veldhuis, Arno-Jan Feddema, Karianne Peterson, Natalia Golender, Marcel Spierenburg, Piet A. van Rijn, Melle Holwerda

**Affiliations:** Royal GD, PO Box 9, 7400 AA Deventer, The Netherlands; University Farm Animal Practice (ULP), Reijerscopse Overgang 1, 3481 LZ Harmelen, The Netherlands; Wageningen Bioveterinary Research (WBVR), Department of Virology, P.O. Box 65, 8200 AB Lelystad, The Netherlands; Kimron Veterinary Institute, Division of Virology, Bet Dagan, PO Box 12, 5025001, Israel; NVWA Incident- and Crisiscentre (NVIC), Netherlands Food and Consumer Product Safety Authority (NVWA), P.O. Box 3511 GG Utrecht, The Netherlands; North-West University, Department of Biochemistry, Centre for Human Metabolomics, North-West University, Potchefstroom, South Africa

**Author notes:** **Article summary line:** Approximately one year after the start of the BTV-3 outbreak in the Netherlands, another serotype, BTV-12, which had never been observed in Europe before, was detected in a sheep sampled on 16 September 2024. This paper describes the first cases, the follow up studies and the first steps to trace the origin of the Dutch BTV-12 outbreak.

**Keywords:** bluetongue, sheep, cattle, serotyping, genotyping, phylogenetic analysis

## Abstract

Bluetongue (BT) is a viral midge borne disease primarily affecting ruminants such as sheep, cattle, and goats. In 2023, the Netherlands reported the first case of bluetongue virus serotype 3 (BTV-3), after being BTV free for eleven years. Since May 2024, vaccination with inactivated BT vaccines for serotype 3 was applied in the Netherlands. Nonetheless, in late June/July 2024, BTV-3 re-emerged and spread over large parts of Europe. In October 2024, BTV-12 was identified by follow-up diagnostics after a BTV-3 vaccinated sheep with signs of BT was tested positive for BTV but negative for serotype 3. This marks a significant event, as BTV-12 had never been reported in Europe. Screening of farms in close proximity to the sheep farm and retrospective analysis of previously BTV PCR positive tested clinical suspicions, resulted in nine BTV-12 affected farms in total. The emergence of BTV-12 in the Netherlands raises important questions about the route of introduction of BT in the Netherlands and mechanisms of viral spread of this specific serotype. Possible adaptation of new BTV serotypes to the European climatic and husbandry conditions prompts reconsideration of surveillance, prevention, and control strategies in relation to changing ecological conditions and vector dynamics. The initial findings, respective studies as well as the initial attempts to trace the origin of BTV-12 are described.

## INTRODUCTION

Bluetongue virus (BTV) is a non-contagious, arthropod-borne virus, which is predominantly transmitted by certain species of *Culicoides* biting midges, so-named competent insect vectors. BTV was historically only present in subtropic and tropic regions between latitudes 35◦S and 50◦N (Gibbs and Greiner, 1994; Zhang et al., 1999). Bluetongue (BT) can remain asymptomatic in ruminants but can also cause (severe) clinical disease and mortality (MacLachlan et al., 2015). The disease can cause significant economic losses in the livestock industry, particularly through morbidity, mortality, and trade restrictions (MacLachlan et al., 2015). Most species of ruminants are susceptible for BTV infection (Niedbalski, 2015). Additionally, infections in new world camelids and incidentally in dogs have been described (Allen et al., 2015; van den Brink et al., 2024). BTV is not zoonotic given that humans are not susceptible for infection (MacLachlan et al., 2015).

The BTV serogroup consists of more than thirty serotypes of which serotypes 1 to 24 are notifiable to the World Organisation of Animal Health (WOAH). Since 1998, several BTV serotypes (1, 2, 3, 4, 6, 8, 9, 11, 16, 25 and 27) have been detected in Europe. In 2006, bluetongue, caused by serotype 8, was discovered in the Netherlands for the first time. From 2009 onwards, BTV was not detected anymore and in 2012, the Netherlands became officially BT-free again (Holwerda et al., 2024). In 2023, an outbreak of a new variant of BTV-3 started in the Netherlands (Holwerda et al., 2024) and has still huge impact on the Dutch sheep and cattle industry (Santman-Berends et al., 2024; van den Brink et al., 2024). In May 2024, three BTV-3 based inactivated vaccines became available aiming to reduce the infection pressure of BTV-3 and reducing clinical signs. Nevertheless, in late June/beginning of July 2024, BTV-3 re-emerged and rapidly spread over large parts of Europe, including Germany, Belgium, Luxembourg, France, Portugal, Spain, Sweden, Norway, Denmark, United Kingdom, Switzerland, Austria and Greece.

Unexpectedly, the first BTV-12 cases in the Netherlands as here described represent a novel virus incursion with potentially significant implications. BTV-12 has previously not been identified in Europe. However, BTV-12 has been recently reported at the European border in Israel (Golender et al., 2023). Its emergence in the Netherlands is notable because its geographical source and route of introduction are unknown. Since this is the second incursion of a novel BTV serotype in Northwestern Europe in a consecutive vector season, it urges the importance to investigate possible viral sources, routes of introduction and active monitoring and surveillance targeting vector borne diseases. In this paper we describe: 1) the first findings like sequencing, virus isolation, and clinical manifestation on a Dutch sheep and dairy cattle farm affected by BTV-12, 2) retrospective screening of samples for BTV-12 submitted in September and October in the Netherlands, and 3) molecular epidemiological study for evaluation of origin of BTV-12 in the Netherlands.

## MATERIALS AND METHODS

### General data on sheep and cattle population in the Netherlands

In 2023, around 1.1 million sheep and 2.6 million dairy cattle (1.6 million ≥2 year and 1 million <2 year) were present on approximately 31,000 sheep farms (of which approximately 10% were professional enterprises (>32 sheep)) and 13,500 dairy cattle farms in the Netherlands. Additionally, there were 14,500 non-dairy cattle herds in the Netherlands (Santman-Berends et al., 2016).

### Notification data on BTV-3 and BTV-12 in the Netherlands in 2024

Notification data was provided by the Netherlands Food and Consumer Product Safety Authority (NVWA, Utrecht the Netherlands), on 20 October 2024. These data included the unique herd number, the date of notification, the involved animal species, the involved bluetongue serotype and the cartographic coördinates, for every clinical BT notification (clinical suspicions and PCR confirmed cases) in 2024. Based on these data a thematic map of the Netherlands was generated with the spmap procedure in Stata 17^®^ presenting the locations of the BTV-3 and BTV-12 notifications in 2024. In the outbreak season of 2023 in the Netherlands, BTV-3 was diagnosed on over 5,000 farms. In 2024, BTV-3 re-occurred and 8,063 farms were affected by BTV until 18 October 2024 (www.nvwa.nl, Figure 1 (grey dots)).

**Figure 1.**
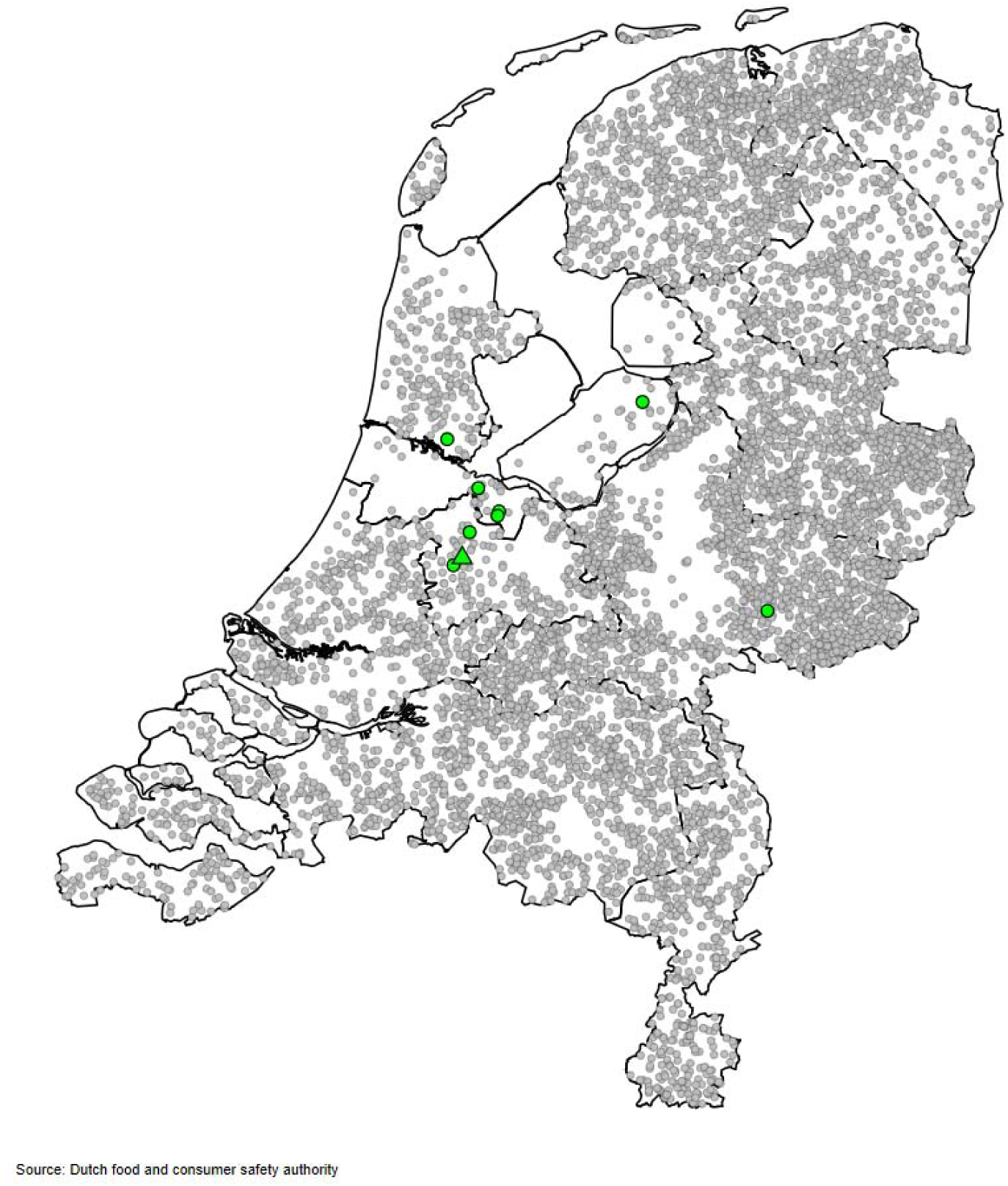
BTV-3 cases notified between July and October 2024 (grey). The green dots represent the location of the nine confirmed BTV-12 cases, the index BTV-12 case is represented by a green triangle.

### Nucleic acid extraction & real-time PCR

To investigate the presence of genomic material of BTV in the collected EDTA-blood, nucleic acids were extracted from 200 µL of EDTA-blood using the Magnapure 96 robotic extraction machine (Roche, Basel, Switzerland) in combination with the Magnapure 96 DNA and viral NA small volume kit. The nucleic acids were eluted in 50 µL. Subsequently, external predenaturation of the double strand RNA of BTV was performed by warming 8 µl of elution in a plate for 1 minute at 99°C and immediately cooled on ice for 5 minutes. Then, 5 µL of the denaturated RNA was added to a new 96-well plate containing 4x TaqMan Fastvirus 1-step Master mix enzyme (Thermo Fischer, Waltham, USA) per well in a total amount of 15 µL with the BTV-specific primers either for a panBTV-approach targeting segment 10 (Hofmann et al., 2008; van Rijn et al., 2012), or to a serotype specific PCR targeting segment 2 of BTV-3 (Maan et al., 2007; Lorusso et al., 2018). The reverse transcription and amplification were performed in Quantstudio 5 (Thermo Fischer) machines using the following real-time PCR program: 50 °C for 10 minutes followed by 95 °C for 1 minute and 45 cycles of 95 °C for 15 seconds and 60 °C for 30 seconds. Analysis was performed using the integrated Quantstudio software.

### Viral genome determination using whole genome sequencing

Sequencing the BTV-12 genome was performed according to the Sequence-Independent-Single-Primer-Amplification-method of Holwerda et al. (2024) and were submitted to the NCBI GenBank database.

### Phylogenetic analyses

The phylogenetic trees were generated using the top thirty hits of the NCBI-data base for each individual segment. Sequences were aligned using MAFFT v7.475 (Katoh and Toh, 2010), phylogeny was reconstructed using maximum likelihood analysis with IQ--TREE software v2.0.3 and 1000 bootstrap replicates (Nguyen et al., 2015) and subsequently visualized using the R package GGTREE (Yu et al., 2017). Bootstrap values above 49 were indicated at the branch nodes and NCBI accession numbers were indicated at the strains.

### Retrospective screening studies of field positive samples for BTV-12

At the day that the causal BTV in the sheep flock was identified as BTV-12/NET2024 (3 October 2024), previously panBTV PCR positive blood samples from BTV-suspected ruminants submitted between 10 and 20 September originating within a radius of twenty kilometers of the sheep case were subject for detailed analysis. After a second case of BTV- 12/NET2024 was identified, all blood samples that were submitted in September and October 2024, to confirm the clinical BT notification, were selected for retrospective analysis. Samples of clinical suspicions which were panBTV PCR positive but BTV-3 PCR negative were subject for further analysis by whole genome sequencing or serotype specific BTV-12 PCR testing.

## RESULTS

### The first index case sheep farm

On 16 September 2024, a blood sample from a seven-month-old ram (Blue Texel) born in the Netherlands, which unexpectedly showed severe clinical signs specific for BT despite vaccination with a BTV-3 vaccine, was submitted to the Dutch national reference laboratory Wageningen Bioveterinary Research (WBVR). The main clinical signs included fever, swollen head, stiffness and lameness, and a tucked up abdomen (Picture 1). After treatment with corticosteroids and NSAID on 14 September, the ram initially improved, but on 18 September, the clinical presentation became more severe resulting in the rams’ death on 20 September 2024.

**Picture 1.**
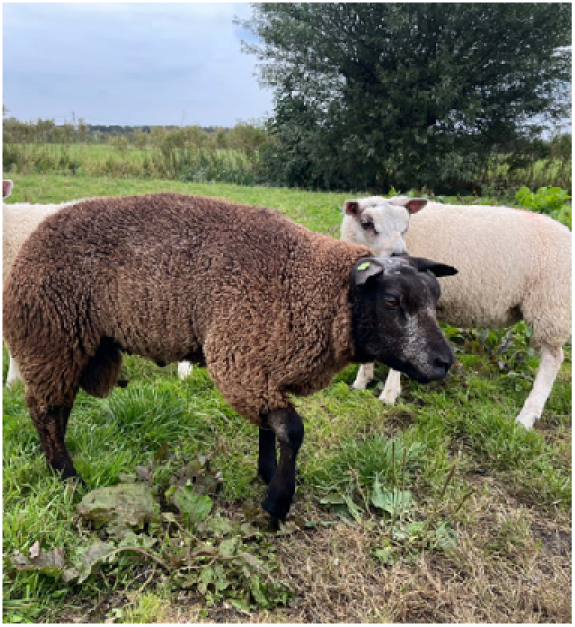
Blue Texel ram showing mild clinical signs of bluetongue on 14 September 2024.

The rams’ blood sample was tested positive twice by panBTV real-time PCR assays (panBTV-PCR), whereas serotype-specific BTV-3 PCR assays tested twice negative. These findings prompted in-depth investigation by whole genome sequencing, according to Holwerda et al., 2024. Genotyping for serotype specific genome segment 2 indicated the presence of BTV serotype 12 in the Netherlands on 3 October 2024. According to the procedure, the blood sample was then sent to the European Union reference laboratory for BT in Madrid (Spain), which subsequently confirmed BTV serotype 12 (BTV-12/NET2024) on 10 October 2024.

The competent authority in the Netherlands, NVWA, was informed and sampled all sheep in the flock the ram originated from. In total, six out of 49 sheep tested panBTV PCR positive and all six were positive for serotype 3. Each of these sheep had shown signs of BT in the previous months but, although not all fully recovered, none of the sheep died. Further analysis using whole genome sequencing showed no co-infections with other BTV serotypes.

The sheep farm, located in the center of the Netherlands (Figure 1, green triangle), had fifty sheep at the time BTV-12/NET2024 was detected. In 2023, between 13 September and the end of November, this farm had been heavily affected by BTV-3/NET2023 showing high morbidity and mortality. In 2024, the entire flock was vaccinated twice with an inactivated BTV-3 vaccine . The second vaccination was administered on 10 August 2024.

### Retrospective studies

A total of nine samples from seven farms were panBTV-PCR confirmed, and one out of nine was negative for serotype 3 (Table 1). This sample was submitted to WBVR on 20 September and originated from a Holstein Friesian heifer on a dairy cattle farm located within five km from the BTV-12 affected sheep farm. Subsequently, seventy out of 73 cattle that were present on this farm, were sampled on 7 October. Three calves (<1 week of age) were excluded from sampling. Except for the initially BTV-12 positive heifer and her calf (date of birth/born on: 13 September 2024), all animals tested negative by the panBTV PCR test. Both the heifer and her calf tested positive in the panBTV PCR and negative in BTV-3 PCR, and revealed BTV serotype 12 based on whole genome sequencing. The initial sample from the heifer was collected on 20 September during a routine farm visit of the local veterinarian. The heifer had calved seven days earlier and, because the heifer had a fever and reddish nostrils, a blood sample was submitted for BTV testing as part of the notification monitoring system. On 7 October, the heifer was clinically recovered, produced according to the farmers’ expectations (>27 kg milk/day), and only showed swollen carpal joints, without signs of lameness (Picture 2a and 2b). Interestingly, her calf showed no clinical signs of BT.

**Table 1.**
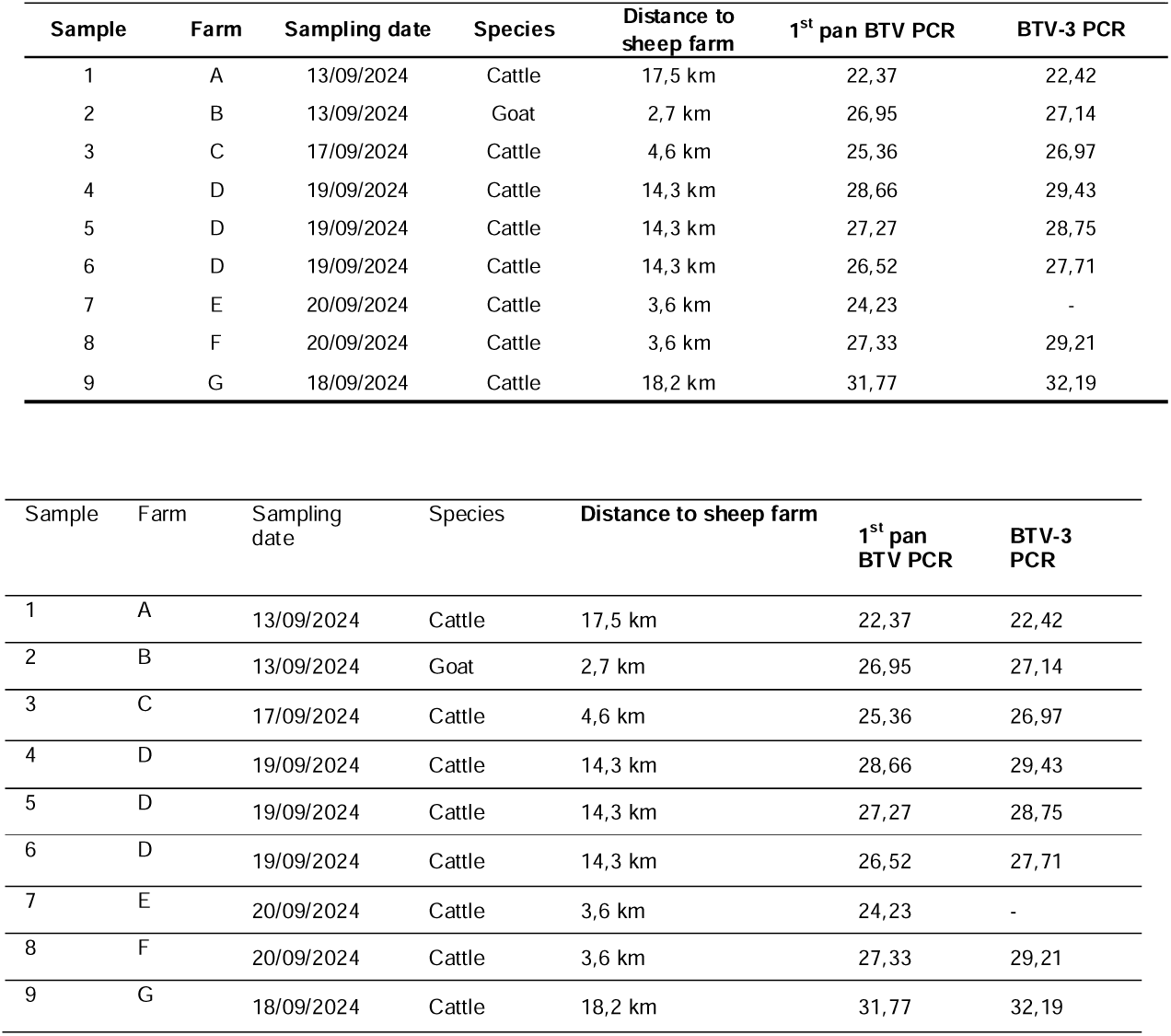
Results of retrospective PCR testing (including cycle threshold values) of samples from the passive surveillance submitted within a radius of twenty km of the sheep farms where BTV-12 was detected.

**Table 2.**
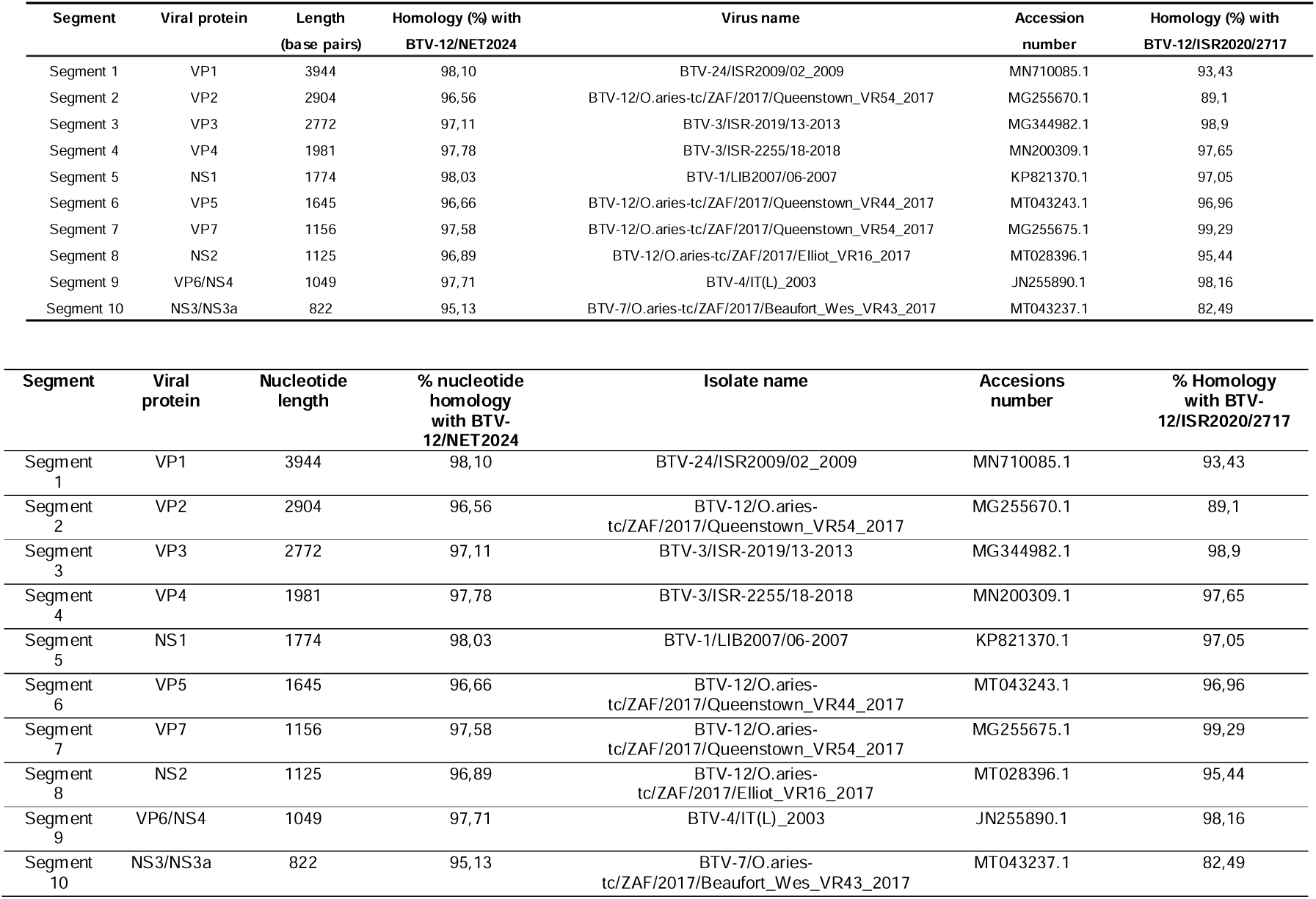
The nucleotide homology of the BTV-12/NET2024 was compared to publicly available sequences from the NCBI database, and to the Israelian BTV-12/ISR2020/2717 virus isolate. The segments of virus isolates with the highest homology are shown. The corresponding NCBI accession numbers of each segment are indicated.

**Picture 2a and 2b.**
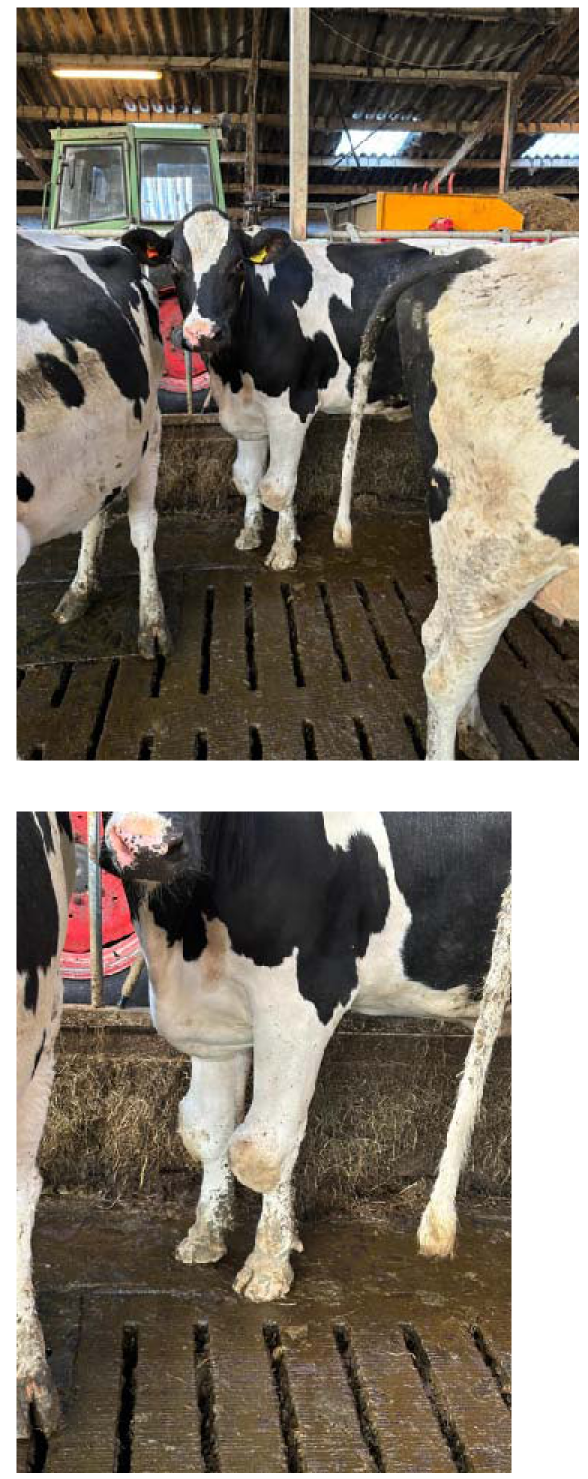
Holstein Friesian heifer showing swollen joints of the carpus at sampling on 7 October 2024.

To investigate to what extend BTV-12 had spread through the Netherlands, all panBTV PCR positive samples that were send in as clinical suspicion towards WBVR in the months September and October of 2024 were tested. A total of 2.519 RNA-samples were subjected to the BTV-3 serotype specific PCR test, and if this test was negative, a BTV-12 specific PCR test was then performed. In total, 2.507 out of 2.519 samples tested positive for BTV-3 (99.52%, 95% C: 99.17-99.75). The remaining twelve samples, submitted between 13 September and 17 October, tested positive for BTV-12. BTV-12 was detected in nine cattle samples from seven cattle farms, and in three sheep samples from two sheep farms. Most of these BTV-12 positive samples originated from the same region as the first two recognized cases. Two farms affected by BTV-12/NET2024 are located at a larger distance from this region (Figure 1; green dots).

### Whole genome sequencing and phylogenetic analysis of BTV-12/NET2024

BTV-12/NET2024 was genetically studied in detail and complete sequences of all ten genome segments were directly recovered from blood samples from the first three described clinical cases. The NCBI accession numbers of the BTV-12/NET 2024 are 4499-4508. Initially, the sequence of genome segment 2 encoding serotype dominant VP2 protein was aligned to sequences available in the NCBI-database (https://www.ncbi.nlm.nih.gov/) to determine the serotype. Preliminary results clearly indicated the highest homology with serotype 12 for segment 2 (Figure 2), but additional phylogenetic analysis was performed for all 10 segments (Figure 3).

**Figure 2.**
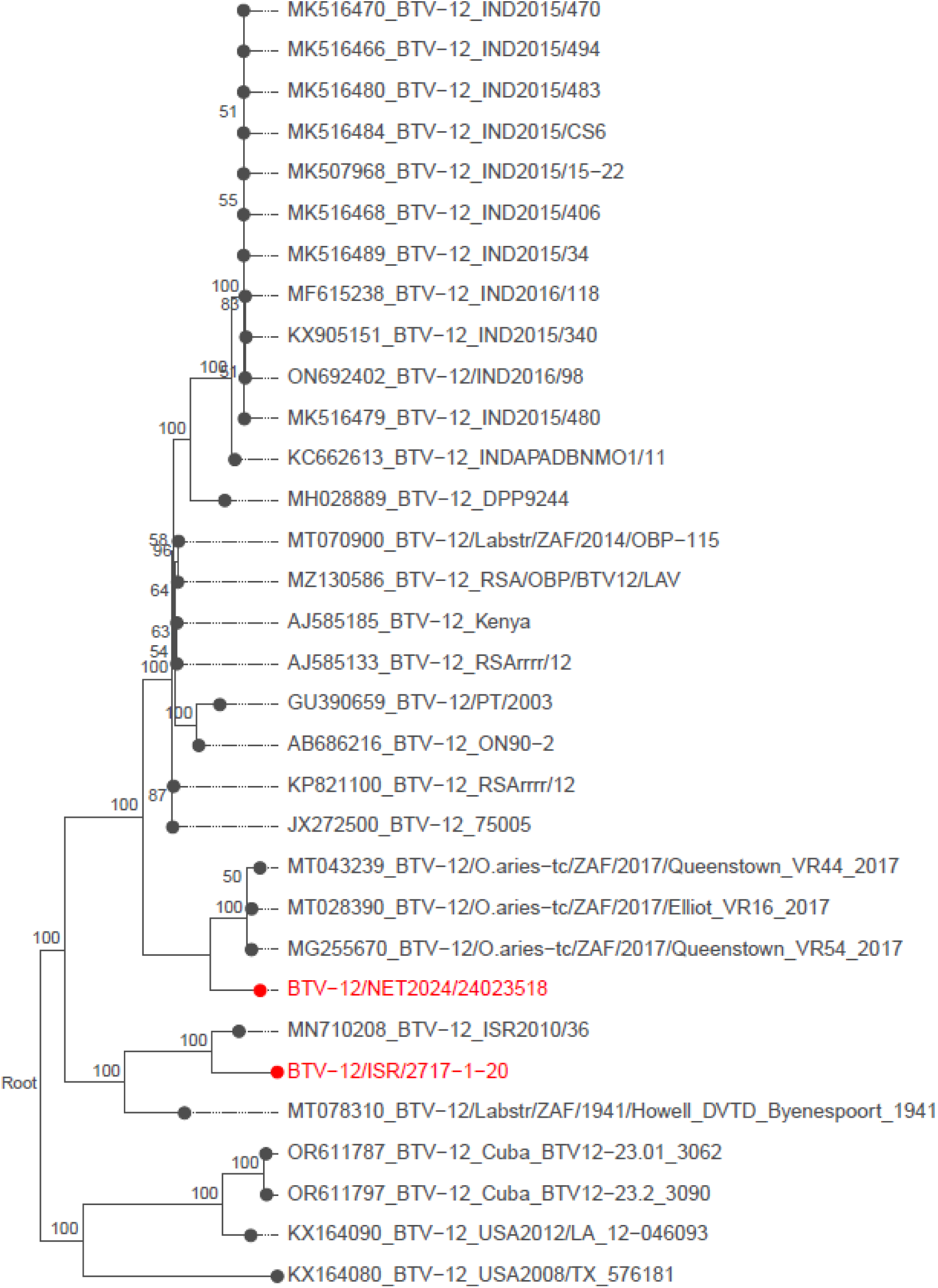
Phylogenetic tree with the relation of segment 2 (VP2) of BTV−12/NET2024/24023518 with the top thirty closest relatives from the NCBI BLAST analysis (accession date Oct 3, 2024) and BTV-12/ISR2020/2717.

**Figure 3.**
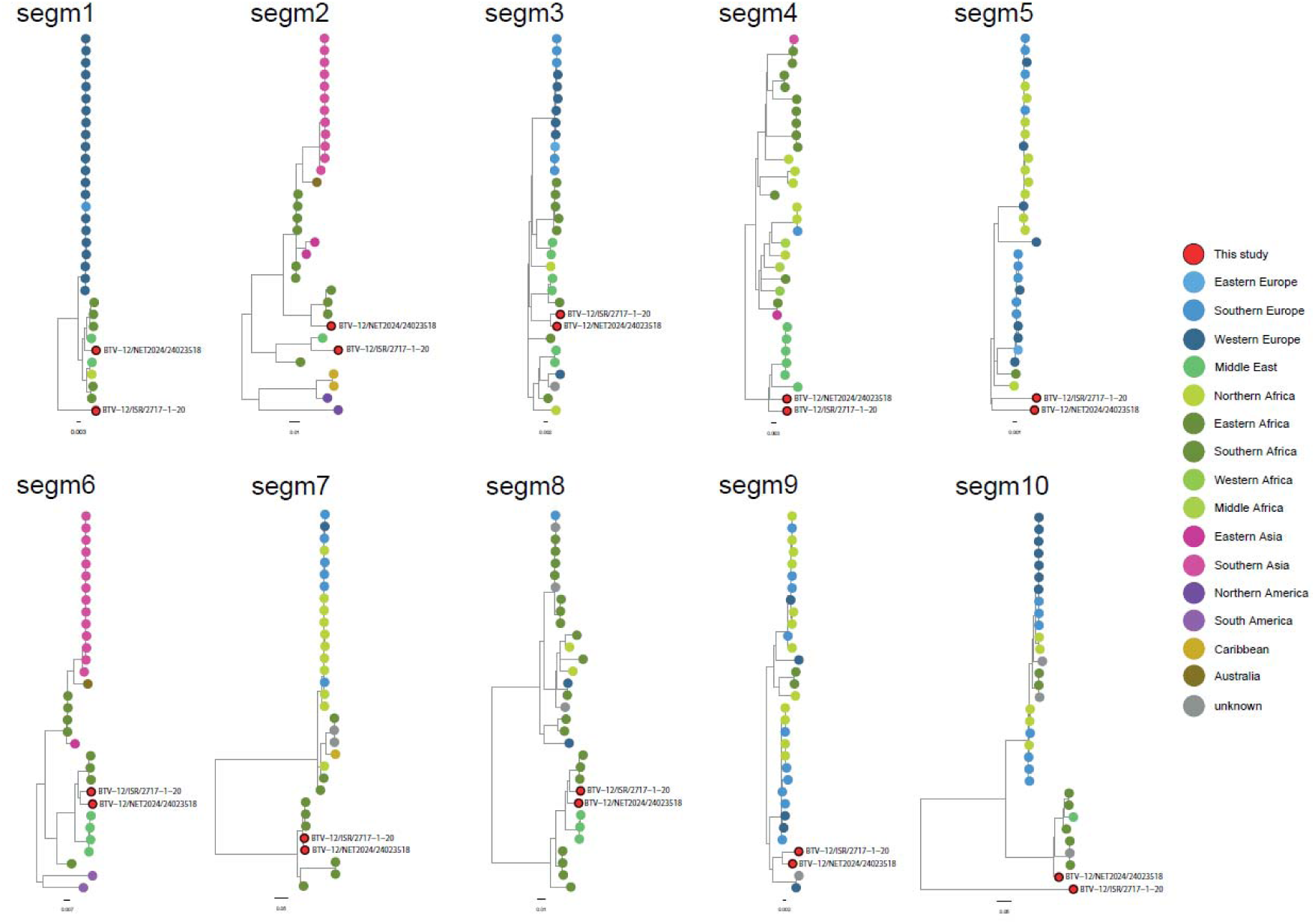
Schematic view of phylogenetic trees for all segments of BTV−12/NET2024/24023518 with the top thirty closest relatives for each segment from the NCBI BLAST analysis (accession date Oct 3, 2024) and BTV-12/ISR2020/2717. Coloured dots represent the origin of the virus isolate. To compare the Dutch (BTV-12/NET2024) and Israeli isolates (BTV- 12/ISR2020/2717), these are highlighted in each tree with a red dot.

## DISCUSSION

The Dutch monitoring and surveillance system is largely based on trust and easy accessible contacts between farmers, veterinarians, Royal GD, WBVR and other veterinary institutes. This system has demonstrated to be highly effective in early detection of emergencies and monitoring trends (Santman-Berends et al., 2016; Dijkstra et al., 2022). In 2024, this monitoring and surveillance system again showed its importance as a Dutch sheep farmer and his veterinary practitioner together with a small ruminant specialist from Royal GD reported clinical signs of BT in a sheep that was prime-boost vaccinated for serotype 3. In a collaborative manner, investigation of this case identified the first European case of BTV serotype 12. Subsequently, BTV-12/NET2024 was retrospectively also found on one additional dairy herd within a radius of five km from the first case. Remarkably, only 1-2 animals on both farms were found positive for BTV-12/NET2024, suggesting that virus spread was either very slow or these were one of the first cases. Further retrospective screening resulted in seven additional BTV-12 affected farms. Most BTV-12 cases were found in the proximity of the initial two cases. Two BTV-12 affected farms are located on a larger distance suggesting transmission by *Culicoides* spp. was unlikely. However, indications of transmission by animal movements were not found. The extremely low number of affected farms and within affected farms and the low number of BTV-12 infected animals could indicate that BTV-12 invaded late in the season of vector activity in Northwestern Europe. A slow virus spread could also be caused by a low vector competence of residential *Culicoides* species for BTV-12/NET2024. Since only a limited number of clinical cases in sheep and cattle were detected in Dutch livestock, it is too early to draw any conclusions regarding the virulence and the potential impact of BTV-12/NET2024 on sheep and cattle. Extended active surveillance is needed to elucidate the prevalence of BTV-12/NET2024 in the Netherlands, since it may also be possible that asymptomatic infections by BTV-12/NET2024 have occurred and possibly remained unnoticed.

In 2008, after the start of massive vaccination against BTV-8/NET2006 in the Netherlands, emerging BTV-6/NET2008 showed a similar phenomenon with only one or two infected animals per farm and some on a larger distance (van Rijn et al., 2012). This BTV-6/NET2008 is strongly related to live-attenuated vaccine (for serotype 6) from South Africa. In the Netherlands, BTV-1 was also detected in an imported animal in 2008 showing the possibility of virus introduction via import of asymptomatic infected animals (van Rijn et al., 2012). Fortunately, BTV-1 and BTV-6 did not massively spread and did not re-emerge in the subsequent year. Illegal use of poorly inactivated BTV-vaccines or infectious BTV as contaminant in other vaccines have been reported and should be considered as a possible virus source (Bumbarov et al., 2016; Ganter, presentation ECSRHM, Turin, 4-5 July 2024). However, two available sequences of segment 2 from live-attenuated vaccine strains for serotype 12 showed low homologies with that of BTV-12/NET2024. Instead, similarities were found with a BTV-12 strain that was identified in Israel at the end of 2020 (Golender et al., 2023), and spread all over the country and the West Bank during the arboviral season of 2021. The unpublished whole genome sequence of BTV-12/ISR2020/2717 was kindly shared and recently submitted to NCBI (accession numbers 0575-0584). Phylogenetic analyses for segments 3 to 9 of BTV-12/NET2024 showed high homology (96-98%) with BTV-12/ISR2020/2717. Nevertheless, homology between BTV-12/2024 and BTV- 12/ISR2020/2717 was much lower for segments 1, 2 and 10 (82-93%). Detailed investigation of segments 1 and 10 showed high homologies with BTV-24/ISR2009/02 (accession number MN710085.1, 98.1% homology) and BTV−3/ISR−2262/2/16 (accession number MG345004.1, 94,15 homology) as detected in the Middle-East. Upon segment 2, the highest homology with segment 2 sequences from Africa suggests that this segment of BTV- 12/NET2024 is closer related to BTV-12 from Africa than from India (Figure 3). Altogether, these results might indicate that seven out of ten segments have been derived from an ancestor circulating in the Middle-East. Further, several reassortment events might have resulted in incorporation of segments 1 and 10 from other co-circulating BTV serotypes, whereas this is unclear for segment 2.

This first detection of BTV-12 in the Netherlands is unique because this serotype had not previously been reported in any European country. The geographical source of the virus remains uncertain, despite the high homology with Israeli sequences for many segments. However, availability of recent sequences is limited. Recent developments are expected to make whole genome sequencing easier and cheaper, in a timely matter and even possible in the field (Sghaier et al., 2022). To enable molecular epidemiological studies aiming at tracing the origin of an emerging virus, it is important to regularly upload sequences with the correct information.

The subsequent/novel incursion in Europe of a second BTV serotype after one year is a significant epidemiological event. Although BTV-3 and BTV-12 are completely different from each other, one common trait is that they were both first observed in the same area of the Netherlands. This raises the question whether there is a relationship between these two introductions or if these events should be regarded as separate occasions. More research should be performed to assess specific factors involved in these BTV incursions in the middle of the Netherlands.

Additionally, these two subsequent BTV incursions raises questions about the future distribution of BTV serotypes and the adequacy of current preventive measures, and signals the need for an expansion of BT and vector surveillance. Monitoring the spread of BTV serotypes is also of critical importance. Historically, BT outbreaks in Europe have followed seasonal patterns corresponding to the activity of *Culicoides* midges, typically peaking in late summer and early autumn. The presence of subsequently new serotypes in the Netherlands may indicate shifting environmental and ecological conditions that facilitate the movement of both vectors and pathogens across geographical boundaries. Given climate change, midge population dynamics may change increasing the risk of novel incursions and spread of BTV serotypes.

Although limited multi-serotype vaccines are available (Savini et al., 2009), current vaccination strategies in Europe are primarily targeting one serotype at the time, such as BTV-1, -3, -4, -8, and -16 by inactivated BT vaccines. It remains the question whether this mono-serotype vaccination approach fulfills the needs in the future to protect ruminant livestock from multiple serotypes of BTV.

Several hypotheses regarding spread of BTV over long distances have been proposed, including the possibility of transport of infected midges by wind, contaminated vaccines or introduction through animal movements or animal products (Mintiens et al., 2008). Here, no proven route of introduction has been yet identified, but as a first step to unravel the introduction of BTV-12, evidence of the geographical origin was found by whole genome sequencing followed by phylogenetic analyses on segment level.

## CONCLUSION

BTV serotype 12 (BTV-12/NET2024) was detected in the Netherlands, after clinical signs of BT were noted in a BTV-3 vaccinated sheep. This was the first report of BTV-12 occurring in the European continent. This prompted a large scale investigation which revealed in total nine BTV-12 positive farms (seven cattle and two sheep farms). Phylogenetic analyses showed high homology with BT-viruses from the Middle-East. Despite this evidence/indication of the geographical origin, the exact virus source, mechanism and route to the Netherlands are currently unknown. This subsequent introduction, after the BTV-3 introduction in 2023, highlights the vulnerability of Europe’s livestock sector to emerging viral threats.

## AUTHOR CONTRIBUTION

RvdB, MvdH, KP were involved in the detection of the sheep case. MvdH was involved in the cattle case. Retrospective investigation of the cattle case was performed by MH, AJF, FH, ME, RV and RG. They were also involved in the laboratory work. MS is involved as official veterinarian from the Dutch Food and Consumer Product Safety Authority (NVWA). RvdB, MH, PvR and IS wrote the initial draft of the manuscript. NG provided epidemiological and molecular data on Israeli BTV-12. All authors did read the draft and gave their comments. RvdB, IS, PvR and MH created the final version of the manuscript and all authors agreed with the content.

## ACKNOWLEDGEMENTS and FUNDING INFORMATION

We would like to thank the colleagues from Royal GD, WBVR and NVWA for their help with the field work and the laboratory investigations. Additionally, we also thank Piet Vellema and Tom McNeilly for their valuable comments on the manuscript. This study is financially supported by the Ministry of Agriculture, Fisheries, Food security and Nature (project number: 1600002757 VZVD).

## ETHICS STATEMENT

The authors confirm that they have adhered to the ethical policies of the journal, as noted on the journal’s author guidelines page. Ethical approval is not applicable.

